# EverGreen: A Calcium Sensing Fluorescent Protein with Ultra-Slow Response Properties

**DOI:** 10.64898/2026.10.29.762894

**Authors:** Ariel Valiente-Gabioud, Oliver Griesbeck

## Abstract

Fluorescent calcium indicators are typically optimized for fast kinetics, but for some bio-engineering projects in synthetic biology slower indicators with long decay time constants may be more practical and easier to apply. We here present a green fluorescent protein calcium biosensor with unusually slow response kinetics. By molecular engineering we were able to extend a slow component of the decay t_1/2_ to 6.1 min. The fluorescence in “EverGreen” could thus be detected for minutes after the underlying calcium fluctuation had already dissipated. This may be useful when constraints in instrumentation and methodology do not allow close coupling of physiological activation of cells and read-out of signals, facilitating directed evolution projects run via plate readers or cell sorters.

## Introduction

Calcium is a ubiquitous messenger involved in a plethora of cellular processes occurring in timescales from milliseconds to hours. As a consequence, many tools were developed to monitor concentrations and fluctuations of calcium in various settings. A prominent role here was taken by fluorescent calcium indicators - both synthetic and genetically encoded - to follow fast fluctuations in cellular free calcium and to decipher spatiotemporal aspects of its regulatory mechanisms^1, 2^. Next to fluorescent indicators, recently another class of fluorescent tools and probes were adding a new facet to the study of cellular calcium. These probes turn elevations of cellular calcium into a more permanent fluorescent record of their signaling history using various molecular principles^3-5^. Thus, a potential read-out of integrated calcium could be performed posthumously at a given time after the actual fluctuations has already ebbed away. In this way, physiological activation, recording and analysis can be disentangled in space and time, giving the experimenter different options for analysis and choice of instrumentation. Here we present a fluorescent probe with properties between the categories of indicators and integrators. It is an intensity-based calcium indicator in which due to unique structural features, Ca^2+^ binding translates very slowly into fluorescent changes, particularly after calcium release. While intriguing on a mechanistic level, the slow allosteric transference renders the biosensor useful for screening projects where larger populations of cells could be probed with microscopy, plate readers or cytometry. In addition, it could be used to filter high frequency calcium transients.

## Results and Discussion

The architecture of the slow-kinetics biosensor is based on GreenT-EC, a low affinity calcium sensor for monitoring calcium in interstitial fluids^6^. It relies on insertion of a calcium sensing moiety derived from Troponin C (TnC)^7^ that was used to replace Tryptophan 148 within the bright green fluorescent protein mNeonGreen^8^ (Fig. 1). The first steps of protein evolution involved a sequential optimization of both linkers at the N- and C-terminus of the inserted calcium binding domain (L1 and L2, respectively) (Fig. 1A). Second, we identified another critical region, comprising amino acid residues C225, R226 and S227 in proximity to L2. Amino acid exchanges within this short region showed a particular potential to impact sensor properties. There were two diversified variants with excellent properties named R1 (CRS -> RTT) and N1 (C225 -> N).

**Figure 1:**
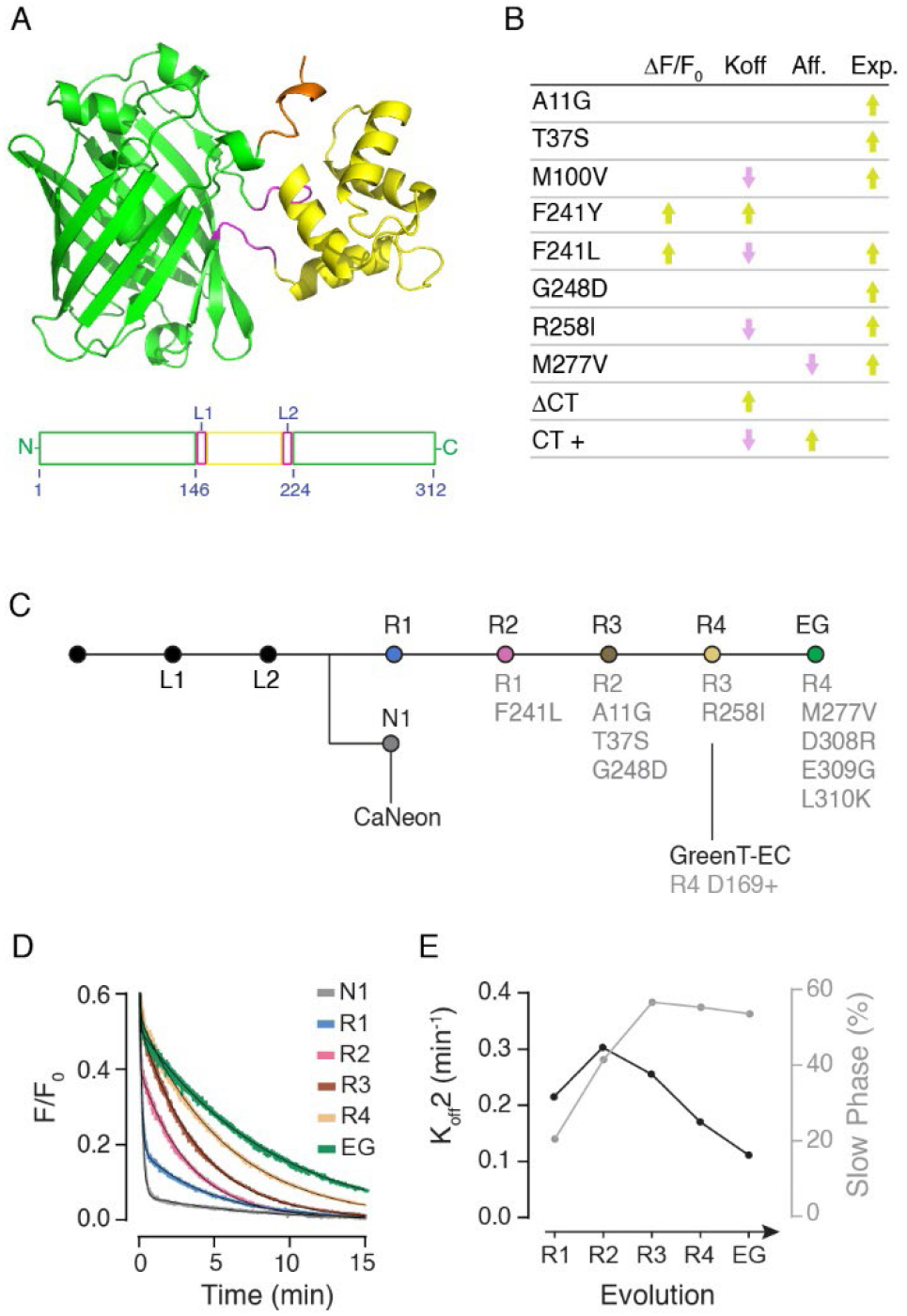
Structure and evolution of EverGreen. **A**) AlphaFold2 model of EverGreen indicating in yellow the calcium binding domain, green for the fluorescent protein, magenta for the linkers and orange the last 9 amino acids of the C-terminus. **B**) Table with a summary of main amino acids affecting the kinetics and expression properties of the sensor. **C**) Evolution of EverGreen. Initial linker optimization is indicated as L1 and L2, followed by the optimization in the region immediately after linker 2 that led to two branches R1 and N1. Further evolutionary steps with single amino acid mutations leading to slow off-rate variants are indicated. **D)** Off-rate curves of variants are overlay where two-phases are evidenced. **E**) The off-rate constant of the slow phase (black line) and the share of the slow second phase to the overall fluorescence decay (grey line) are plotted.

During the process of protein evolution, we closely monitored the fluorescence change in response to calcium binding, the affinity and off-rate properties of a large number of variants. Several single mutations led to significant changes in both affinity and kinetics. They also presented significant changes in response and expression/folding efficiency (Fig. 1B). In Fig. 1C we schematize the sequence of diversification steps that finally led to EverGreen (EG). While both variants R1 and N1 presented similar responses, they showed significantly different kinetics behaviors. While the decay kinetics of N1 was mono-exponential, R1 was brighter in the bound state and revealed a two-phase decay. We chose R1 for further engineering and directing the decay time of the indicator into an ultra-slow regime. Variant N1 (Fig. 1C) turned out to be the starting point for generating a high affinity cytosolic calcium biosensor^9^.

One particular amino acid residue, phenylalanine 241 appeared to be very instrumental for manipulating expression, fluorescence changes and binding dynamics. Within the structure of the sensor, this residue is pointing inwards towards the chromophore. Another amino acid that showed a strong impact in protein expression levels and off-rate kinetics in vitro was M100. Mutations of M100 however abolished calcium responses in mammalian cells, thus it was not followed up further. The position of these amino acids suggests that the strong effect on response kinetics is due to a change in the process of chromophore deprotonation upon calcium dissociation and not on the binding process of calcium itself. In particular, the mutation F241L, introduced to generate variant R2 (Fig. 1C), presented a two-component double exponential decay, with a dominant fast-phase t^1/2^ of 1.5 sec (59 % share) and a second, smaller but much slower component of t^1/2^ of 2.3 min (41% share). Another mutation, F241Y, increased the decay rate 2-fold. We reasoned that the slow afterglow of the fluorescence would endow the probe with integrator-like properties that could be put to use in certain experimental situations. By a combination of directed evolution and rational design we thus intended to improve the properties of the protein, aiming at optimizing fluorescence response, expression and folding, while enhancing the secondary slow decay component to allow several minutes of detectable fluorescence afterglow for subsequent post-experimental analysis.

We then identified critical amino acid residues that determine protein folding and expression properties by means of error prone mutagenesis, and further used site saturation in those specific locations (A11, T37, G248, R258, M277). The variant R3 (Fig. 1C) contained the mutations A11G, T37S and G248D and was used as template further due to its exquisite expression properties. We found that the mutation R258I (variant R4) drastically reduced the off-rate and was therefore included in the final variant. In addition, M277V increased the overall fluorescence change and was included as well. Over all generations we were able to increase the share of the slow component to the overall fluorescence decay of the sensor. Finally, when performing structural modeling of the sensor using I-Tasser, we observed that the last 9 amino acids at the C-terminal end established interactions with the calcium binding minimal TnC domain. We observed that removing the C-terminus completely had a pronounced effect, reducing the decay time two-fold. Truncation thus accelerated decay, which suggested that the same residues could be engineered to exert the opposite effect of slowing the decay further. Accordingly, we rationally inserting a series of positively charged amino acid variants to the C-terminal tail of the sensor which allowed us to identify combinations of residue exchanges that prolonged the slow phase further and increased its share in the overall decay. Replacing the original, predominantly acidic C-terminal pentapeptide DELYK with more basic stretches (e.g. RGKK, RNKYK, RGKYK, RRKYK) progressively slowed fluorescence decay, while acidic replacements accelerated it. The final variant, “EverGreen”, had a major absorbance peak maximum in the calcium bound state at 504 nm. In the absence of calcium, minor absorbance peaks at 350 nm and 400 nm were apparent (Fig. 2A). Its maximal fluorescence change was 88.6-fold in vitro (Fig. 2C) upon calcium addition, with a high specificity against magnesium (Fig. 2C). The excitation and emission maxima were 504 nm and 515 nm, respectively. The fluorescence decay after rapid calcium chelation was well described by two exponentials, with k_off_1 = 26 min^−1^ (τ = 2.3 s), carrying 46% of the fluorescence response, and k_off_2 = 0.11 min^−1^ (τ = 8.8 min), carrying the remaining 54% (Fig. 2D). The engineering steps that slowed fluorescence decay also slowed down the apparent association rate. The on rate was at 0.054 s−1, clearly set to low-pass filter signals within cells (Fig. 2E, F). In vitro affinity titrations of the sensor revealed a Kd of 234 nM, making the sensor well placed for recording events within the cytosol of living cells. The main spectroscopic properties of EverGreen *in vitro* are summarized in Table 1.

**Table 1:** Spectroscopic properties of recombinant EverGreen. ΔF/F_0_, fluorescence change upon calcium binding; EC (mM^-1^ cm^-1^), extinction coefficient of the bound state; K_D_ (nM), dissociation constant; QY, quantum yield of the bound state; Brightness (Br.) is obtained as QY*EC; n_H,_ Hill coefficient; Koff1 and Koff2 (min^-1^), fast and slow off-rate kinetics respectively of the two-phase dissociation curves. The percentage of the response obtained during the slow second phase is indicated; Kobs (s^-1^) is the observed on-rate kinetics calculated at 235 nM free calcium concentration. Errors were calculated from the fitting and technical replicates.

| | $\Delta F/F_0$ | EC | QY | Br. | $k_D$ | $n_H$ | $K_{off1}$ | $K_{off2}$ | $K_{obs1}$ | pKa |
| --- | --- | --- | --- | --- | --- | --- | --- | --- | --- | --- |
| Mean | 88.6 | 96.6 | 0.76 | 72.9 | 234 | 0.94 | 25.7<br>(46%) | 0.1136<br>(54 %) | 0.0536 | 6.32 |
| SD | 4.1 | 3.5 | 0.03 | 3.9 | 7 | 0.02 | 0.1 | 0.0001 | 0.0004 | 0.03 |

**Figure 2:**
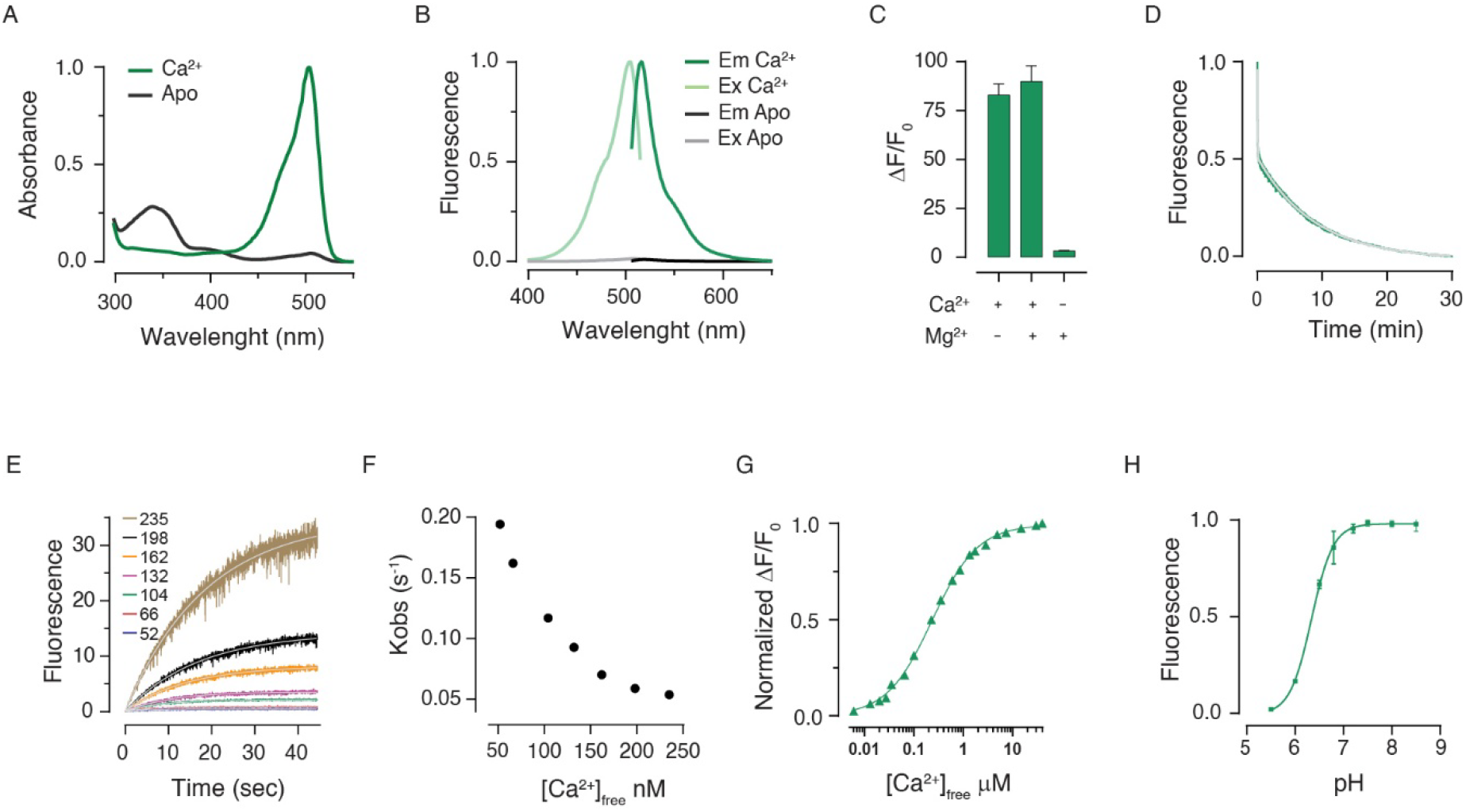
Spectroscopic characterization of recombinant EverGreen. **A**) Absorbance changes in the absence (grey) and presence (green) of calcium. The major absorbance peak maximum was at 504 nm. Minor absorbance peaks at 350 nm and 400 nm in the absence of calcium corresponded to dark forms of the chromophore. **B**) Excitation and emission spectrum at 0 µM (gray) and 200 µM (green) Ca^2+^. The excitation and emission maxima are 504 nm and 515 nm, respectively. **C**) Maximal fluorescence changes of EverGreen in response to calcium and/or magnesium (+ is 1 mM, - is 0 mM). D) Off-rate kinetic of EverGreen. The protein was prepared in MOPS 30 mM, KCl 100 mM, Mg^2+^ 2.5 mM, Ca^2+^ 200 µM and rapidly mixed with BAPTA 200 mM. **E**) On-rate kinetics of EverGreen was measured by rapidly mixing solutions of protein in MOPS 30 mM, KCl 100 mM, Mg^2+^ 1 mM with MOPS 30 mM, KCl 100 mM, Mg^2+^ 1 mM containing increasing concentration of Ca^2+^. A single exponential was used for fitting single association kinetics and the obtained Kobs value was plotted against the calcium concentration. **F)** A decrease in the Kobs is observed at increasing concentrations of calcium. These results suggest a mechanism by which there is an equilibrium between at least two species of the sensor, and only one can progress to a fluorescent species upon calcium binding. **G)** In vitro affinity titration curve. **H**) pKa of the protein obtained by preparing pH solutions in MOPS/MES containing 100 mM Ca^2+^.

We next set out to characterize EverGreen in mammalian cells (Fig. 3). As basic fluorescence at resting calcium levels was extremely dim, we fused EverGreen to a red reference protein, called mCyRFP^10^. By choosing appropriate linkers we ensured that the fusion did not result in any detectable FRET. The reference protein allowed identifying transfected cells, while green fluorescence would only develop after stimulation or activation of cells. In addition, EverGreen signals could be plotted as green/red ratios, if desired. We compared its properties to that of the green calcium indicator GCaMP6f^11^. In HeLa cells stimulated with 200 µM histamine GCaMP6f reported a relatively steep onset transient that peaked about 30s after bath application and was not detectable over baseline anymore 5-10 min later (Fig. 3A-C). EverGreen signals, in contrast, rose slowly, peaking at 1.5 minutes, and then decayed over tens of minutes, still reporting about 73% of its peak response at 10 min, 62% at 15 min and 45% at 25 min (Fig 3A-C). We then also quantified EverGreen behavior by flow cytometry using HEK293 cells, using ionomycin to raise cytosolic calcium and EGTA to chelate it (Fig. 3D, E). Also here, GCaMP6f signals peaked fast but also decayed fast after calcium chelation using EGTA. EverGreen signals, in contrast, decayed slowly but still persisted detectably, at 2 min (47% of peak signal) and 10 minutes (11 % of peak signal) after removing calcium.

**Figure 3:**
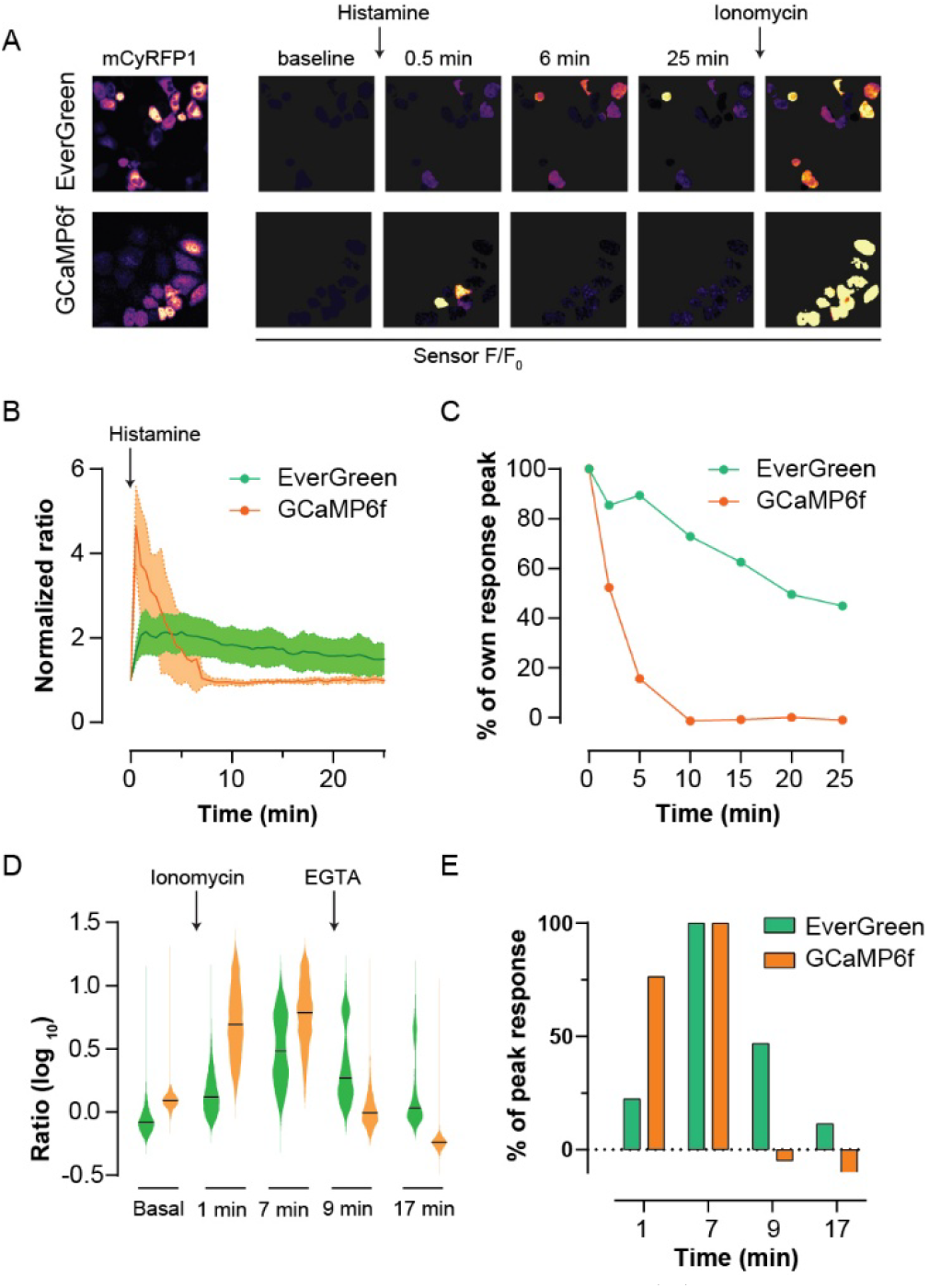
EverGreen in Mammalian Cells. (**A**) Representative images of HeLa cells expressing EverGreen (top) or GCaMP6f (bottom) fused to mCyRFP1: mCyRFP1 reference (left) and sensor (green channel) fluorescence normalized to the pre-stimulus frame (F/F_0_), shown for mCyRFP1-positive cells. Application of histamine (200 µM) and Ionomycin/CaCl_2_ (2.5 µM/0.5 mM) is indicated. EverGreen rises more slowly and is still elevated 25 min later. Scale bar, 50 µm. (**B**) Ratio traces normalized to baseline; lines are medians, shaded bands the interquartile range (n = 22 EverGreen and 24 GCaMP6f cells, two independent experiments). (**C**) The same data expressed as percentage of each sensor’s peak response. (**D**) Flow cytometry of HEK293 cells after treatment with ionomycin and subsequent wash with EGTA. Violins are single cell green/red ratios for gated cells; bars indicate the median. (**E**) Median response at each time point as percentage of the indicator’s peak response.

In summary, we here present a biosensor that takes its place between conventional fast kinetics fluorescent calcium indicators and integrators that sum up the history of activations and turn it into an irreversible fluorescence signal. We performed extensive protein engineering to ensure a robust fluorescence signal upon activation and generated fusions with a red reference protein to enable ratioing responses. The slow decay time was found to be due to some peculiar molecular features and mechanisms. The off-rate was bi-phasic, with a fast and a slow component. The phases could be clearly separated, and particularly the slow-phase could be enhanced by molecular engineering. Calcium dissociation from TnC typically occurs orders of magnitude faster than what we observed in the slow decay phase of EverGreen. Presumably slow chromophore relaxation is the rate-limiting step here, not calcium dissociation. The absorbance spectrum of EverGreen is different from that of calcium biosensors such as GCaMP6. In GCaMPs, the absorbances corresponding to the neutral and anionic chromophore change in concert after binding calcium. In EverGreen, there is an additional absorbance band peaking around 350 nm (Fig. 2A), which mainly exchanges with the anionic peak absorbance, reminiscent of the photo-switchable protein Dreiklang, where the 350 nm species was assigned to a hydrated form of the chromophore^12^.

The lack of an irreversible fluorescence of the sensor still requires the read-out to take place within minutes of the stimulation experiment, which could be seen as a disadvantage. However, this also allows for repeated cycles of stimulation and decay taking place, without having to discard cells and experiments after a single trial. Possibly further engineering can be applied to slow the decay time down further or, finally, result in variants with a potentially irreversible fluorescence as in previous integrators.

The presence of an absorbance band at 350 nm under calcium-free conditions, next to the barely detectable peak of the protonated species at 400 nm, is reminiscent of the photo-switchable fluorescent protein Dreiklang^12^. Potentially, EverGreen can be further developed into a photo-switchable sensor with integrator-like properties. The current variant, however, barely shows detectable photo-switching (data not shown). Another drawback of the current version may be the relatively slow on-rate which results in pronounced low-pass properties that favor detection of slower stimuli versus rapidly fluctuating signals.

Despite these limitations, the first implementation of EverGreen provides detection technology for calcium with new and interesting features. It may find its use in experiments where large populations of cells need to be monitored quickly for on-off stimulus activation and allows further passaging and probing collected cells with additional conditions. This could facilitate a number of experiments in directed evolution and synthetic biology.

## Methods

### Molecular biology

The vector pRSET-B (Invitrogen) was used for library construction, bacterial screening and protein production, and pcDNA 3.1+ (Invitrogen) for expression in mammalian cells. Constructs were assembled by homology-based SLiCE cloning. Random mutagenesis was performed by error-prone PCR with the GeneMorph II kit (Agilent), and standard PCR and site-directed mutagenesis with Herculase II Fusion DNA polymerase (Agilent). To obtain a ratiometric readout, mCyRFP1^10^ was fused to the N-terminus of EverGreen and, for comparison, of GCaMP6f, through a 20-aa hydrophobic flexible linker^13^ chosen to prevent FRET between the two proteins. Primers were purchased from Eurofins Genomics, and all constructs were verified by Sanger sequencing.

### Protein expression and purification

His-tagged proteins were produced in E. coli BL21 (Invitrogen) by overnight growth at 37 °C in 50 ml auto-induction medium (LB with 0.05% (w/v) D-glucose, 0.2% (w/v) lactose and 0.6% (v/v) glycerol). Cells were pelleted (6,000 × g, 10 min, 4 °C) and resuspended in 10 ml resuspension buffer (20 mM Na_2_PO_4_, 300 mM NaCl, 20 mM imidazole) containing protease inhibitors (4 µM PMSF, 20 µg/ml pepstatin A, 4 µg/ml leupeptin), 5 µg/ml DNase and 10 µg/ml RNase (all Sigma-Aldrich). Cells were lysed by sonication on ice (7 min; Bandelin Sonoplus, duty cycle 0.8, power 0.8), and the lysate was cleared by centrifugation (20,000 × g, 30 min, 4 °C). The supernatant was incubated with 150 µl of a 6% (v/v) Ni-IDA agarose suspension (Jena Bioscience) for 1 h at 4 °C under gentle agitation. The beads were transferred to 1 ml gravity-flow columns (Qiagen) and washed with 10 ml resuspension buffer, and the protein was eluted in 700 µl elution buffer (20 mM Na_2_PO_4_, 300 mM NaCl, 300 mM imidazole) and dialyzed against MOPS buffer (30 mM MOPS, 100 mM KCl, pH 7.2).

### Spectroscopy

Absorbance spectra were recorded on a Cary 300 UV-Vis spectrophotometer (Varian), and excitation and emission spectra on a Cary Eclipse fluorescence spectrophotometer (Varian). Proteins were measured in MOPS buffer supplemented with 0.4 mM EGTA (Ca^2+^-free) or 0.2 mM Ca^2+^ and 1 mM Mg^2+^ (Ca^2+^-saturated). The fluorescence change (ΔF/F0) was measured in 96-well plates in a Tecan plate reader. F0 was recorded in MOPS buffer with 0.4 mM EGTA, and the Ca^2+^-bound fluorescence in MOPS buffer with 1 mM Ca^2+^, in the absence or presence of 1 mM Mg^2+^. Selectivity was assessed by measuring the fluorescence in 1 mM Mg^2+^ without Ca^2+^. The molar extinction coefficient (ε) was calculated from the absorbance of the alkali-denatured chromophore at 443-446 nm (ε = 44 mM^−1^ cm^−1^). Proteins were prepared in MOPS buffer supplemented with 0.2 mM Ca^2+^, and the absorbance spectrum was recorded before and after addition of NaOH to a final concentration of 0.1 M. mNeonGreen was measured in parallel as a reference (ε = 114 mM^−1^ cm^−1^ in our hands).

The quantum yield was determined relative to mNeonGreen (quantum yield 0.80^8^) with the slope method. Absorption and emission spectra of three serial 1:2 dilutions (starting at an absorbance of 0.1) were acquired in the same cuvette, the integrated emission was plotted against the absorbance at the excitation wavelength, and the slope was compared with that of mNeonGreen. Brightness was calculated as ε × quantum yield.

To determine the pKa, a series of MOPS/MES-buffered solutions supplemented with 100 mM Ca^2+^ was prepared, with pH adjusted from 5.5 to 8.5 using NaOH and HCl. Triplicates of 200 µl of buffer containing 0.5-1 µM protein were prepared in a 96-well plate for each pH value and incubated for 10 min at room temperature, and the emission spectra were recorded. The relative fluorescence at the emission maximum was plotted against pH and a sigmoidal curve was fitted to obtain the pKa.

Kinetic measurements were done on a Cary Eclipse fluorescence spectrophotometer (Varian) fitted with an RX pneumatic drive unit (Applied Photophysics), at room temperature (about 23 °C), with excitation at 480 nm and emission at 520 nm. For the macroscopic off-rate (Koff), a Ca^2+^-saturated protein solution (30 mM MOPS, 100 mM KCl, 2.5 mM MgCl_2_, 200 µM CaCl_2_, about 0.2-1 µM protein, pH 7.2) was rapidly mixed with a BAPTA solution (30 mM MOPS, 100 mM KCl, 20 mM BAPTA, pH 7.2) at an injection pressure of 3.5 bar. Traces were recorded at 0.1 s intervals for 30 min and fitted with a double-exponential function in Prism, giving Koff1, Koff2 and the share of each phase. For the macroscopic on-rate (Kobs), Ca^2+^-free protein (30 mM MOPS, 100 mM KCl, 1 mM MgCl_2_, about 0.2-1 µM protein, pH 7.2) was rapidly mixed with the same buffer containing increasing Ca^2+^ concentrations, giving final free Ca^2+^ concentrations of 39-628 nM. Traces were acquired at 12.5 ms intervals for 45 s and fitted with a single exponential. Free Ca^2+^ concentrations were calculated with the WEBMAXC Standard server.

The Ca^2+^ affinity was determined in MOPS buffer (30 mM MOPS, 100 mM KCl, pH 7.2) containing 10 mM EGTA, 1 mM Mg^2+^ and increasing concentrations of Ca^2+^, as described previously^14^. Free Ca^2+^ concentrations were calculated with WEBMAXC Standard, and fluorescence was measured on a Cary Eclipse spectrometer. The dissociation constant (KD) and the Hill coefficient (nH) were obtained by plotting ΔF/F0, normalized to its maximum, against the log10 of the free Ca^2+^ concentration (in mol/l) and fitting a sigmoidal curve with variable slope in Prism.

### Bacterial colony screening

Libraries were transformed into E. coli XL1-Blue cells and plated on LB-agar with ampicillin (Roth), giving 500-800 colonies per plate; depending on library size, 20-50 plates were screened per round. After overnight incubation at 37 °C, the plates were kept at 4 °C for 24 h to favor protein maturation before imaging. A home-built screening platform was used to image the plates before and after addition of 10 ml of screening buffer (30 mM MOPS, 100 mM KCl, 50 mM CaCl_2_, 0.5% agar, pH 7.2) kept at 42 °C, and the images were analyzed with a customized Python routine^6^. After each round, selected clones were expressed and purified and characterized by their response to Ca^2+^, extinction coefficient, affinity and kinetics, and the best candidates were tested in mammalian cells. GCaMP6s was used as control indicator for calibration and comparison.

### Cell lines

HeLa and HEK 293T cells (Invitrogen) were grown in high-glucose Dulbecco’s modified Eagle medium with pyruvate (Gibco) supplemented with 10% fetal bovine serum (Gibco), L-glutamine, penicillin and streptomycin at 37 °C and 5% CO_2_.

HeLa cells were seeded in 35 mm glass-bottom dishes (MatTek) and transfected one day before imaging with the mCyRFP1-EverGreen or mCyRFP1-GCaMP6f constructs using Lipofectamine 3000 according to the manufacturer’s instructions. Imaging was done in Hank’s balanced salt solution (HBSS) supplemented with 1 mM MgCl_2_ and 3 mM CaCl_2_ on a Leica STELLARIS 5 confocal microscope. After recording a baseline, cells were stimulated with 200 µM histamine and, at the end of the recording (27.5 min), with 2.5 µM ionomycin. Images were processed in ImageJ with custom macros. Noise was reduced with a median or Gaussian filter (2 pixels), and saturated and background pixels were excluded by thresholding: the original image was divided by the binary mask so that excluded pixels were assigned N/A, which prevents zero values from biasing the ROI averages. The processed green (EverGreen or GCaMP6f) and mCyRFP1 channels were then divided to obtain a ratio image, ROIs were drawn on individual cells, and the mean ratio of each ROI was exported and normalized to its baseline (mean of the frames before histamine).

### Flow cytometry

HEK 293 cells were seeded in 35 mm dishes (Corning) and transfected with the mCyRFP1-EverGreen or mCyRFP1-GCaMP6f constructs using Lipofectamine 3000. After 24h cells were detached with Versene (Gibco), collected in HBSS supplemented with 3 mM CaCl_2_ and pelleted at 1,000 × g for 5 min. Cells were resuspended in 1.5 ml MOPS buffer supplemented with 3 mM CaCl_2_ and 1 mM MgCl_2_ and transferred to 5 ml round-bottom tubes with cell-strainer cap (Corning). Samples were measured in an Attune NxT acoustic focusing cytometer (Thermo Fisher Scientific) before treatment, 1 and 7 min after addition of 2.5 uM ionomycin, and 2 and 10 min after subsequent addition of 4 mM EGTA. At least 100,000 events were recorded per condition and analyzed with FlowJo v10. Cells were gated on forward and side scatter, and doublets were excluded. The BL1-A channel (488 nm excitation, 530/30 nm emission) was used for EverGreen and GCaMP6f, and the YL2-A channel (561 nm excitation, 620/15 nm emission) for mCyRFP1. For each cell the green/red ratio was calculated; the median ratio of each sample was used to express the response as the percentage of the peak response of each indicator (basal = 0%, maximum after ionomycin = 100%). Data were plotted with GraphPad Prism.

## Supporting Information

The amino acid sequence of EverGreen: MVSKGEEDNMGSLPATHELHIFGSINGVDFDMVGQGSGNPNDGYEELNLKSTKGDLQFSPWILVPHIGYGFHQYLPYPDGMSPFQAAMVDGSGYQ VHRTMQFEDGASLTVNYRYTYEGSHIKGEAQVKGTGFPADGPVMTNSLTAADLGWDSEEELSECFRIFDFDGNGFIDREEFGDIIRLTGEQLTDEDVDEIFGDSDTDKNGRIDFDEFLKMVENVQLTDNRTTKKTYPNDKTIISTLKWSYTTDNGKRYRSTAITTYTFAKPMAANYLKNQPVYVFRKTELKHSKTELNFKEWQKAFTDVMGMRGKYK

## AUTHOR INFORMATION

## Author Contributions

A.V. generated the sensor, performed directed evolution, in vitro spectroscopy, microscopy and analysis via cytometry. A.V. and

O.G. designed the research, wrote and edited the paper.

## Notes

The authors declare no competing financial interests.

## ACKNOWLEDGMENT

We would like to thank Anja Moritz for technical assistance.

## REFERENCES

(1) R.Y. Tsien; Intracellular signal transduction in four dimensions: from molecular design to physiology. Am J Physiol. 1992, 263, 723–8.

(2) Boulware & Marchant; Timing in cellular Ca^2+^ signaling. Curr. Biol. 2010, 18, R769–R776.

(3) Fosque, B. F.; et al. Labeling of Active Neural Circuits in Vivo with Designed Calcium Integrators. Science 2015, 347, 755–760.

(4) Kim C.K.; et al. A Molecular Calcium Integrator Reveals a Striatal Cell Type Driving Aversion. Cell 2020, 183, 2003-2019.e16.

(5) Huppertz M.-C., et al. Recording physiological history of cells with chemical labeling. Science 2024, 383, 890–897.

(6) Valiente-Gabioud, A. A.; et al. Fluorescent sensors for imaging of interstitial calcium. Nat. Commun. 2023, 14, 6220.

(7) Thestrup, T.; et al. Optimized Ratiometric Calcium Sensors for Functional in Vivo Imaging of Neurons and T Lymphocytes. Nat. Methods 2014, 11, 175–182.

(8) Shaner, N. C.; et al. A Bright Monomeric Green Fluorescent Protein Derived from Branchiostoma lanceolatum. Nat. Methods 2013, 10, 407– 409.

(9) Nevelchuk S.; et al. Morphotype-specific calcium signaling in human microglia. J. Neuroinflammation 2024, 21, 175.

(10) Laviv T.; et al. Simultaneous dual-color fluorescence lifetime imaging with novel red-shifted fluorescent proteins. Nat. Methods 2016, 13, 989–992.

(11) Chen, T.-W.; et al. Ultrasensitive Fluorescent Proteins for Imaging Neuronal Activity. Nature 2013, 499, 295–300.

(12) Brakemann, T.; et al. A Reversibly Photoswitchable GFP-like Protein with Fluorescence Excitation Decoupled from Switching. Nat. Biotechnol. 2011, 29, 942–947.

(13) Arslan, M.; et al. Effect of non-repetitive linker on in vitro and in vivo properties of an anti-VEGF scFv. Sci. Rep. 2022, 12, 5485.

(14) Tsien, R. Y., & Pozzan, T. Measurement of cytosolic free Ca? with quin2. Methods in Enzymology (1989), 172, 230–262.

